# Photopharmacological activation of adenosine A_1_ receptor signaling suppresses seizures in a mouse model for temporal lobe epilepsy

**DOI:** 10.1101/2024.07.04.602052

**Authors:** Jeroen Spanoghe, Paul Boon, Marijke Vergaelen, Maren De Colvenaer, Tina Mariman, Kristl Vonck, Evelien Carrette, Wytse Wadman, Erine Craey, Lars E. Larsen, Mathieu Sprengers, Jeroen Missinne, Serge Van Calenbergh, Dimitri De Bundel, Ilse Smolders, Robrecht Raedt

## Abstract

Up to 30% of epilepsy patients suffer from drug-resistant epilepsy (DRE). The search for innovative therapies is therefore important to close the existing treatment gap in these patients. The adenosinergic system possesses potent anticonvulsive effects, mainly through the adenosine A_1_ receptor (A_1_R). However, clinical application of A_1_R agonists is hindered by severe systemic side effects. To achieve local modulation of A_1_Rs, we employed a photopharmacological approach using a caged version of the A_1_R agonist N^6^-cyclopentyladenosine, termed cCPA. We performed the first *in vivo* study with intracerebroventricularly (ICV) administered cCPA to investigate the potential to uncage sufficient amounts of cCPA in the hippocampus by local illumination in order to suppress hippocampal excitability and seizures in mice. Using hippocampal evoked potential recordings, we showed a reduction in hippocampal neurotransmission after photo-uncaging of cCPA, similar to that obtained with ICV injection of CPA. Furthermore, in the intrahippocampal kainic acid mouse model for DRE, photo-uncaging of CPA in the epileptic hippocampus resulted in a strong suppression of seizures. Finally, we demonstrated that intrahippocampal photo-uncaging of CPA resulted in less impairment of motor performance in the rotarod test compared to ICV administration of CPA. These results provide a proof of concept for photopharmacological A_1_R modulation as an effective precision treatment for DRE.

## 1. Introduction

Of the estimated 50 million epilepsy patients worldwide, more than one third suffers from drug-resistant epilepsy (DRE)^1–3^. In these patients, currently available antiseizure medication (ASM) fails to achieve seizure freedom or causes unacceptable side effects. In the search for innovative therapies to close the existing treatment gap in epilepsy patients, adenosine receptor agonists have been the subject of many preclinical studies^4–11^. The adenosinergic system plays an important endogenous anticonvulsive role in the central nervous system (CNS). Extracellular adenosine concentrations have been shown to rise during seizures^12–14^, which is hypothesized to be an important factor for the spontaneous termination of seizures. These seizure-suppressive effects of adenosine can mainly be attributed to binding of adenosine to the adenosine A_1_ receptor (A_1_R) subtype. Activation of this G_i_-coupled receptor leads to a suppression of neurotransmitter release through inhibition of voltage-gated Ca^2+^ channels and a decrease in neuronal excitability by opening of K^+^ channels, causing membrane hyperpolarization^15^. Some of the highest expression levels of A_1_Rs are found in the hippocampus^16^, from which seizures often originate in case of temporal lobe epilepsy (TLE). As one of the most common forms of DRE^17^, TLE therefore marks itself as an ideal target for adenosine-based interventions.

Administration of adenosine or A_1_R agonists has been demonstrated to be effective in models for DRE^18,19^. However, clinical translation of such therapies is hindered by severe side effects of systemic administration, such as bradycardia and hypothermia, due to the ubiquitous expression of A_1_Rs throughout the body. To take advantage of the strong seizure-suppressive effects of the A_1_R, strategies are required that allow for local activation of these receptors at the site of the seizure focus. In the past, antiseizure effects have been achieved with chronic local delivery of adenosine via infusion with osmotic minipumps, implantation of adenosine-releasing polymers and genetically-engineered adenosine-releasing cells^8,20–24^. These strategies all involve a local but invariable, continuous increase of adenosine concentrations. We are now investigating a novel approach that has the potential to deliver a more controlled modulation of A_1_Rs, namely photopharmacology.

Through a combination of photochemistry and pharmacology, the field of photopharmacology involves the use of light to provide high spatiotemporal control over the activity of drugs^25–27^. With the technique of “photocaging”, a photoremovable protective group (or “photocage”) is covalently bonded to a bioactive ligand, thereby rendering it inactive. Activity can be restored by breaking the photosensitive bond using illumination of a specific wavelength, a process called “photo-uncaging”. Photopharmacology has become an emerging field of research and in recent years its therapeutic potential is increasingly being explored *in vivo*. For instance, photocaged compounds have been tested for control over metabotropic glutamate type 5 and opioid receptors in analgesia studies^28–30^, the adenosine A_2A_ receptor in Parkinson’s disease^31^ and the adenosine A_3_ receptor in psoriasis^32^. Yet, so far this technique has only seen limited application in epilepsy research. A first proof of concept for its therapeutic potential was delivered by Yang *et al.,* who used local uncaging of caged γ-aminobutyric acid to terminate paroxysmal activity induced by 4-aminopyridine in the neocortex of rats *in vitro* and *in vivo*^33,34^.

For photopharmacological modulation of A_1_R activity, we recently caged the A_1_R agonist N^6^-cyclopentyladenosine (CPA) with the coumarin-derivative 7-(diethylamino-4-(hydroxymethyl)-2H-chromen-2-one (DEACM), which is sensitive to light of 405nm wavelength^35^ (**Figure 1**). This caged CPA (cCPA) has previously been validated *ex vivo* in our laboratory and displays a 1000-fold reduced affinity for the A_1_R compared to CPA^36^. In acute hippocampal brain slices, positioned on a multielectrode array (MEA) and superfused with 3 µM cCPA, illumination with millisecond flashes of 405 nm light at an intensity of 4 mW uncaged an estimated concentration of 30 nM CPA. This was sufficient to suppress stimulation evoked potentials (EPs), which reflects a reduction in neurotransmission and neuronal excitability and thus indicated successful A_1_R activation with cCPA. Furthermore, this approach was also capable of suppressing epileptiform bursts in the high-potassium hippocampal slice model^37^.

**Figure 1.**
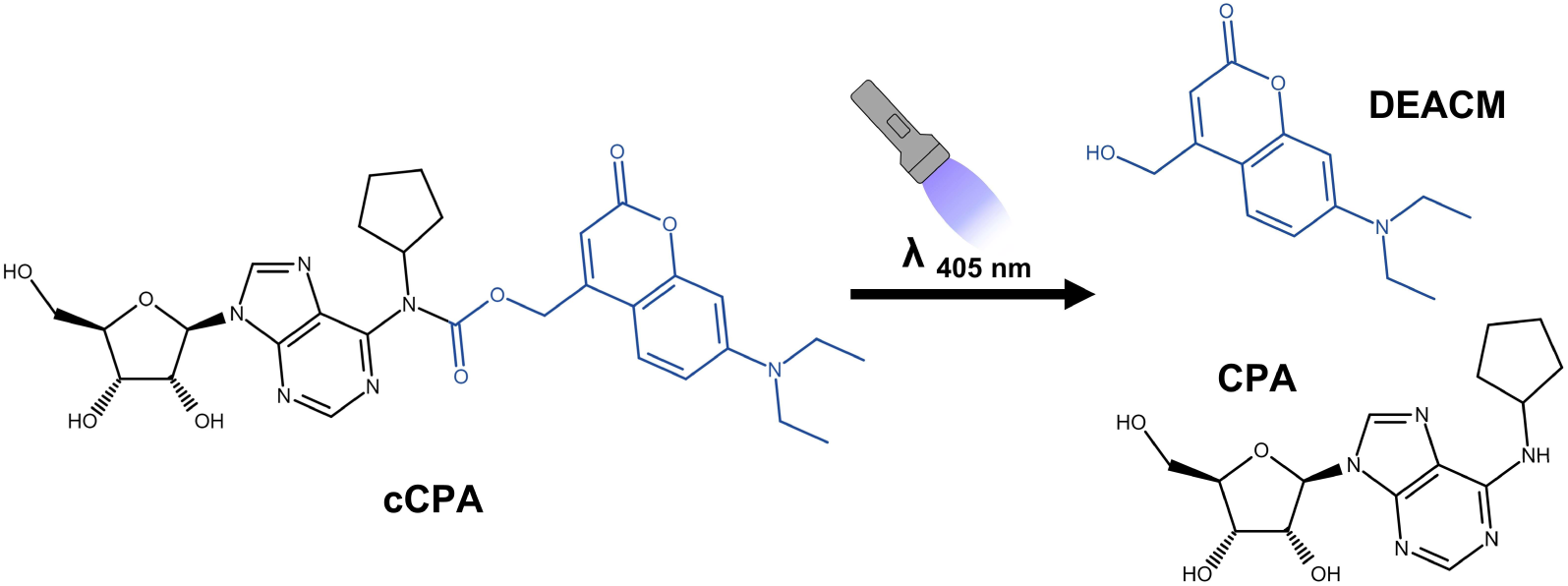
Structure of caged CPA (cCPA) and schematic representation of its uncaging into CPA and DEACM after exposure to 405 nm illumination. CPA: N^6^-cyclopentyladenosine; DEACM: 7-(diethylamino-4-(hydroxymethyl)-2H-chromen-2-one.

In the present study, we explored the use of cCPA for the first time *in vivo*. First, we investigated if intracerebroventricularly (ICV) administered cCPA could be uncaged using local illumination through an implanted optical fiber in the hippocampus of anesthetized mice. We studied effects on hippocampal EPs similar to our previous *ex vivo* study, where a reduction in EPs reflects A_1_R activation. These hippocampal EPs were evoked by electrical stimulation of presynaptic axons of the perforant path (PP), which causes depolarization of postsynaptic granule cells in the dentate gyrus (DG) resulting in a field excitatory postsynaptic potential (fEPSP)^38^. Synchronous generation of action potentials in the granule cells induces a “population spike (PS)”, which is represented as a negative deflection on the fEPSP (**Figure 2**). Next, we investigated whether local A_1_R activation with this photopharmacological approach can be used to suppress seizures in the intrahippocampal kainic acid (IHKA) mouse model, which displays chronic recurrent hippocampal seizures that are resistant to several frequently used ASMs^39,40^. To this end, effects on seizure frequency were examined after ICV administration of cCPA combined with local hippocampal illumination. Finally, we investigated possible effects of our photopharmacological treatment on locomotor activity, as administration of A_1_R agonists is known to cause sedation^41,42^. Since we aim to achieve a more spatially controlled activation of A_1_Rs with cCPA in order to avoid such side effects, we compared the effects between CPA administration and uncaging of cCPA on the performance of mice on the accelerating rotarod test.

**Figure 2.**
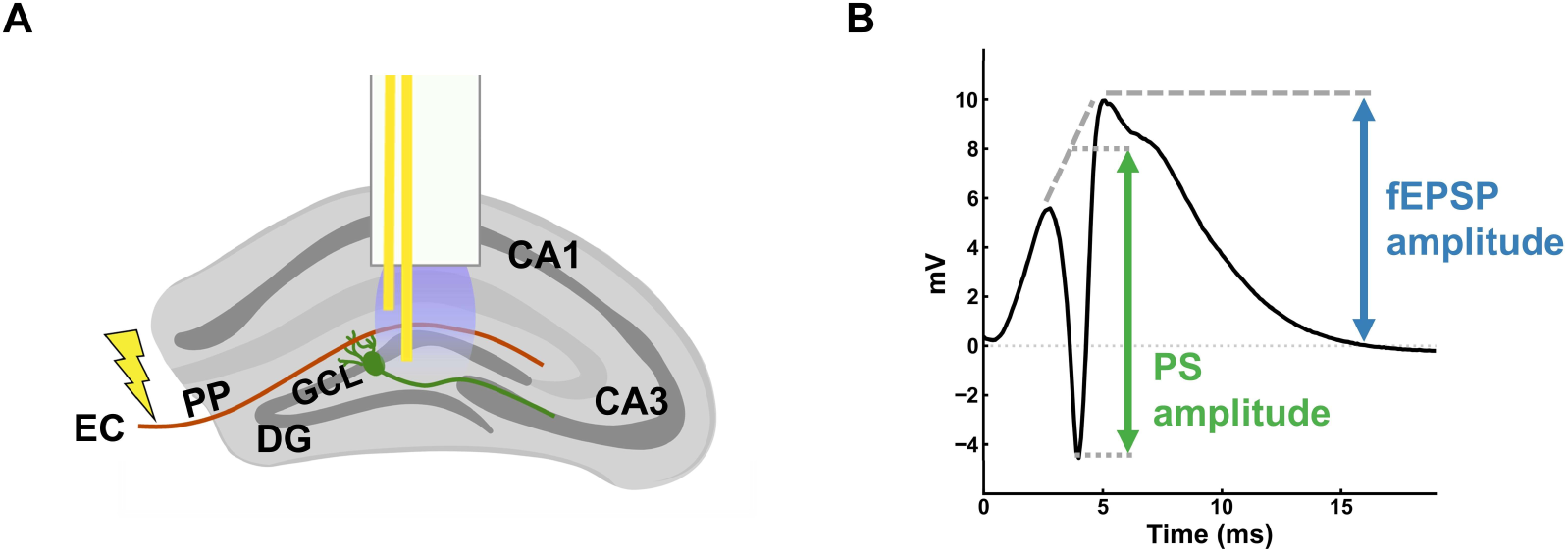
(**A**) Illustration of the hippocampus with an optrode implanted for evoked potential (EP) recording. Electrical stimulation of the perforant path elicits EPs in the granule cell layer of the dentate gyrus. (**B**) Example of a hippocampal EP constituting of a positive field excitatory post-synaptic potential (fEPSP) and a negative population spike (PS). CA: cornu ammonis; DG: dentate gyrus; EC: entorhinal cortex; GCL: granule cell layer; PP: perforant path.

## 2. Material & Methods

### 2.1. Chemicals

The A_1_R agonist N^6^-cyclopentyladenosine (CPA) was acquired from Tocris Bioscience (Bristol, UK). The DEACM used as photocage was acquired from Sigma-Aldrich (Saint Louis, USA). Photocaged CPA (cCPA) was synthesized as previously reported^36^. Dimethylsulfoxide (DMSO, Tocris Bioscience) was used as solvent for both CPA and cCPA. Solutions containing cCPA were prepared in the dark and shielded from ambient light during all experiments.

### 2.2. Animals

Experiments were performed in adult male C57Bl/6J mice (8 weeks old at the start of experiments), obtained from Envigo (The Netherlands). Animals were housed under environmentally controlled conditions (temperature 21-22°C, relative humidity 40-60%) at a fixed 12-hour light/dark cycle. Food and water were available *ad libitum*. Prior to the start of all experiments, all animals were group housed. Animals were housed individually after injections and implantations. All procedures were conducted in accordance to the European Directive 2010/63/EU and were approved by the Animal Experimental Ethical Committee of Ghent University (ECD 20/21).

### 2.3. Evoked potential recordings with CPA & cCPA administration

#### 2.3.1. Stereotaxic surgeries

In a terminal experiment, healthy mice were anesthetized with isoflurane (5% at 2 l/min for induction, 2% at 0.5 l/min for maintenance) and mounted into a stereotaxic frame (Stoelting, USA). Body temperature was monitored and kept stable using a rectal probe connected to a heating pad. A midline incision was made in the scalp to expose the skull and holes were drilled for intracerebroventricular (ICV) injection and placement of electrodes. A 5 µl Hamilton Neuros syringe (Hamilton Company, USA) was inserted in the right lateral ventricle (coordinates: −0.3 mm AP and +1.0 mm ML relative to bregma, −2.2 mm DV relative to dura). On the contralateral side, a bipolar recording electrode (200 µm tip separation) was lowered into the DG (coordinates: −2.0 mm AP and +1.3 mm ML relative to bregma, −2.0 mm DV relative to dura) and a bipolar stimulation electrode (500 µm tip separation) into the perforant path (coordinates: −3.7 mm AP and +2.0 mm ML relative to bregma, −1.5 mm DV relative to dura). Bipolar electrodes were fabricated from two twisted polyimide-coated stainless-steel wires of 70 µm (recording) and 120 µm (stimulation) diameter (California Fine Wire, USA). An epidural screw electrode (stainless steel, 1.57 mm diameter, Bilaney Consultants, Germany) was placed in the frontal bone as ground/reference electrode. For experiments with illumination, the recording electrode was combined with a multi-mode optical fiber (400 µm core diameter, 0.50 NA, FP400URT, Thorlabs, Germany) into an optrode, the distal contact of the bipolar electrode protruding 300 µm from the tip of the optical fiber. Prior to insertion into the brain, light output was measured at the tip of the optical fiber using a photodiode power sensor (S120VC coupled to PM100A, Thorlabs, Germany) and set to the desired intensity. The position of the recording and stimulation electrodes was adjusted using electrophysiological feedback until optimal evoked potential (EP) waveforms were acquired. Recordings were started once all electrodes and the syringe were set in place. After conclusion of the EP recordings, animals were euthanized with an overdose of sodium pentobarbital (1500 mg/kg, i.p.).

#### 2.3.2. Recording protocol

Recording of hippocampal field potentials was done using a custom MATLAB-based script (MathWorks, USA), controlling a USB-6211 NI-DAQ card (National Instruments, USA) for data acquisition and stimulation. Field potentials were high-pass filtered at 0.1 Hz, amplified 248 times and digitized at 10 kHz with a 16-bit resolution (input range of ±10 V). Recordings were stored on a PC for off-line analysis. Every 10 seconds, a 6-second sweep of electroencephalography (EEG) was recorded. To record EPs, the perforant path was stimulated at the start of every sweep by delivering biphasic square-wave pulses (200 µs per phase) through the stimulation electrode, generated by a constant current linear stimulus isolator (Digitimer, UK). Input-output (I/O) curves were constructed by stimulating every 10 seconds at increasing current intensities (50-500 µA in increments of 50 µA, 500-1000 µA in increments of 100 µA), repeated four times. The intensity evoking about 50% of the maximal PS amplitude on the averaged I/O curve was determined and used as stimulation intensity for the subsequent EP recordings.

In a first batch of animals, the effect of two ICV administered CPA doses was studied. After at least 10 minutes of baseline EP recording, a 5 µl solution was injected in the lateral ventricle at a rate of 10 µl/min with the Hamilton syringe controlled by a Quintessential Stereotaxic Injector (Stoelting, USA). Animals received either 0.25 µg CPA (n = 5), 1.25 µg CPA (n = 7) or vehicle (2.5% DMSO in 5 µl saline, n = 4). Recordings were continued for at least 60 minutes after injection.

In a second batch of animals, cCPA was administered ICV either with (cCPA-light, n = 8) or without illumination (cCPA-dark, n = 7) of the dorsal DG. In the cCPA-light group, the optrode implanted in the hippocampus was connected to a 405 nm light-emitting diode (M405FP1, Thorlabs, Germany) via a patch cable (M98L01, Thorlabs, Germany). The illumination protocol consisted of 100 ms light pulses at a frequency of 0.1 Hz (1% duty cycle) with an intensity of 8 mW (63.5 mW/mm² at optical fiber tip). After baseline EP recordings, 25 µg cCPA (dissolved in 1.25 µl DMSO) was injected ICV at a rate of 10 µl/min. Recordings were continued for at least 60 minutes, with a 20-minute illumination period applied 10 minutes after injection. Directly after conclusion of the recording, the optrode was removed from the brain and its light output was measured. In 2 out of 8 mice, the output had decreased by more than 50% of the starting value (due to damage to the fiber or coverage of optical fiber tip by blood) and thus recordings were excluded. In the cCPA-dark group, baseline EP recordings were followed by injection of 25 µg cCPA and at least 60 minutes of recording after injection without illumination.

### 2.4. cCPA administration in the IHKA model

#### 2.4.1. IHKA injections

Mice (n = 12) received unilateral hippocampal injections of kainic acid (KA) for the induction of status epilepticus. Animals were anesthetized and placed into the stereotaxic frame as described above (section 2.3.1). After exposing the skull, a hole was drilled for injection of KA into the right hippocampus (coordinates: −2.0 mm AP and +1.5 mm ML relative to bregma). A glass capillary loaded with KA solution (4 mg/ml KA in saline) was lowered into the dorsal hippocampus (−1.8 mm DV relative to dura) and 50 nl was infused at a rate of 100 nl/min using a Nanoliter injector system (World Precision Instruments, USA). The capillary was left in place for 5 minutes after injection and then slowly retracted to prevent backflow. Afterwards, the skin was closed with non-degradable stitches and animals were observed after waking up from anesthesia for signs of status epilepticus.

#### 2.4.2. Implantations

Three weeks after KA injection, mice underwent a second stereotaxic surgery for implantation of an optrode in the right hippocampus and a guide cannula above the left lateral ventricle. The optrodes consisted of a bipolar electrode (polyimide-coated stainless-steel wire, California Fine Wire, USA) combined with a multi-mode optical fiber (400 µm core diameter, 0.50 NA, FP400URT, Thorlabs, Germany) and had an average light transmission of 82 ± 2% compared to the output measured at the tip of the patch cable. Animals were anesthetized and placed into the stereotaxic frame as described above (section 2.3.1). Holes were drilled for insertion of an optrode in the injected hippocampus (coordinates: −2.0 mm AP and +1.5 mm ML relative to bregma, −2.0 mm DV relative to dura) and implantation of a guide cannula (C316G-SPC, Bilaney Consultants, Germany) above the lateral ventricle on the contralateral site (coordinates: −0.3 mm AP and +1.0 mm ML relative to bregma, −0.8 mm DV relative to dura). Additional burr holes were made for the placement of 2 epidural screw electrodes and 2 anchor screws. One epidural electrode was placed over parietal cortex contralateral to the optrode for recording of cortical EEG, the other epidural electrode was placed over frontal cortex to serve as ground/reference electrode. Electrode leads were attached to a connector and assembled into a headcap together with the optrode and guide cannula using dental acrylic cement.

Mice were allowed to recover from surgery for at least one week. Following recovery, they were handled and habituated to the procedure of ICV injection, which consisted of one researcher restraining the awake animal and immobilizing the head while a second researcher inserted a Hamilton syringe through the guide cannula to manually inject vehicle solution (1.25 µl DMSO) in the ventricle (−2.2 mm DV relative to dura). The syringe was kept in place for 1 minute after which it was slowly retracted to prevent backflow.

#### 2.4.3. EEG recordings in the IHKA model with cCPA administration

After implantation and habituation, mice were connected to an EEG setup via their headcap. This setup consisted of a unity gain preamplifier, a 6-channel cable, a commutator and 512x amplifier, leading to a USB-6259 NI-DAQ card (National Instruments, USA). Signals were high-pass filtered at 0.15 Hz and digitized at 2 kHz (16-bit resolution, input range ±10 V). Recordings were controlled via custom made MATLAB (MathWorks, USA) software and stored on a PC for off-line analysis. Baseline EEG was recorded for 4 hours and evaluated for the presence of electrographic hippocampal seizures. Animals that displayed clear seizures were selected for a cross-over study (7 out of 12 animals).

The cross-over design consisted of 3 treatments: ICV administration of 25 µg cCPA without illumination (cCPA-dark), 25 µg cCPA with illumination (cCPA-light) and vehicle (1.25 µl DMSO) with illumination (vehicle-light). Animals were randomly assigned to 1 of 3 sequences to receive all 3 treatments, with a 2-day washout in between each treatment. On the day of the treatment, animals were connected to the EEG setup and at least 2 hours of baseline EEG was recorded. Animals were disconnected for ICV injection and afterwards immediately returned to the setup to continue EEG recording for at least 4 hours. For treatments including illumination, animals were also connected with the implanted optrode to a 405 nm laser diode (LP405-MF300, Thorlabs, Germany) via a patch cable (M127L01, Thorlabs, Germany). Illumination was performed for a period of 2 hours immediately after injection and consisted of 100 ms light pulses with an intensity of 50 mW (396.8 mW/mm², output measured at the patch cable tip) at a frequency of 0.1 Hz. Six out of 7 animals underwent all 3 treatments, one animal lost its headcap after the first treatment with cCPA-dark. Of those 6, the EEG of one animal during vehicle treatment could not be analyzed due to a bad connection to the EEG setup.

In a subset of animals (n = 3), additional recordings were performed to control for possible effects of the photocage DEACM, which is released upon uncaging of cCPA. After recording baseline EEG for at least 2 hours, animals were injected with DEACM solution (10.14 µg in 1.25 µl DMSO), equimolar to the cCPA solution used, after which EEG recordings were continued for at least 4 hours.

### 2.5. Behavioral testing

Healthy mice (n = 6) were implanted with an optrode and a guide cannula and habituated to ICV injections following the same procedures as described above (section 2.4.2).

The effects of CPA and cCPA administration on locomotor activity were tested with the accelerating rotarod test^43^. Mice were placed on a rotarod apparatus (Mouse RotaRod, Ugo Basile, Italy), set to accelerate from 4 rpm to 40 rpm over 300 seconds and the time to the end of each trial was recorded. The trial ended when the animal either fell off the rotarod, completed 2 full rotations when clinging on to the rod or when it ran for 300 seconds. Animals first underwent training for 3 consecutive days, consisting of 10 rotarod trials separated by a 3-minute rest period each day. This allowed the animals to learn the task and reach a plateau in their performance by the third day of training^44^. After completion of the training, animals were tested on the rotarod after administration of 4 different treatments in the following order: ICV administration of vehicle (1.25 µl DMSO) with illumination, 25 µg cCPA without illumination (cCPA-dark), 25 µg cCPA with illumination (cCPA-light) and CPA (1 µg) with illumination. Each treatment was separated by 2 days of wash-out.

On the day of treatment, animals were connected to the EEG setup via their headcap and one hour of baseline EEG was recorded. Afterwards, animals were disconnected for ICV injection and immediately returned to the setup to continue EEG recording for 30 minutes. For treatments including illumination, animals were also connected to the 405 nm laser diode and illumination (100 ms pulses at 0.1 Hz, 50 mW) was performed during the 30 minutes recording after injection. After conclusion of these recordings, animals were disconnected and placed on the rotarod to perform the accelerating rotarod test.

### 2.6. Histological verification

The correct placement of the guide cannula and optrode was histologically verified in implanted animals after conclusion of the experiments (**Supplementary Fig 1**). Mice were anesthetized and 0.5 µl of Evans Blue dye solution (1% w/v in saline) was injected through the guide canula prior to euthanasia with an overdose of sodium pentobarbital (1500 mg/kg, i.p.). Transcardial perfusions were performed with phosphate-buffered saline followed by 4% paraformaldehyde solution. Brains were isolated, cryoprotected with 30% sucrose solution and snap-frozen in isopentane. Coronal sections of 40 µm thickness were made on a cryostat (CM1950, Leica, Germany) and stained with cresyl violet (0.1% w/v).

### 2.7. Data analysis

#### 2.7.1. Evoked potentials

Hippocampal EPs were analyzed using custom MATLAB (MathWorks, USA) software, measuring the field excitatory post-synaptic potential (fEPSP) amplitude and population spike (PS) amplitude for each evoked response. The fEPSP amplitude was calculated as the maximum value of the fEPSP peak. The PS amplitude was calculated as the distance between the negative peak of the PS and the line connecting the positive peaks before and after the PS (**Fig 2 B**). Values were normalized to the mean of baseline and averaged per 5 minutes.

#### 2.7.2. EEG power

Power spectral analysis of all recorded EEG data was performed using custom scripts in Python (v3.8.5, Python Software Foundation). The differential EEG signals derived from the bipolar electrode in the hippocampus were high-pass filtered at 1 Hz (1^st^-order Butterworth) and segmented into 1-second epochs with 50% overlap. These segments were windowed (Blackmann-Harris) after which the Fast Fourier algorithm was used to compute the power spectra, divided over the following frequency bands: total power (1-100 Hz), delta (1-3 Hz), theta (4-12 Hz), beta (13-30 Hz), gamma (31-100 Hz). For subsequent analyses, frequencies around 50 Hz (48-52 Hz) were excluded to avoid power line interference. Power values were log-transformed into decibels (10*log10), the average power per Hz was taken for each frequency band and the difference in power to baseline was calculated.

For the EP recordings (section 2.3), power spectra were obtained from the last 5 seconds of each 6-second sweep and averaged per 5 minutes. From 1 recording in the high dose group, EEG power could not be analyzed due to incomplete EEG-sweeps. For the EEG recordings in the IHKA model (section 2.4), the average power spectra were calculated from 5 interictal 1-minute segments which were randomly selected from the 2-hour baseline period and from the 2-hour period after injection. For the EEG recordings from the rotarod test (section 2.5), power spectra were calculated from the last 20 minutes of the baseline recording and the 20-minute segment from minute 10 to minute 30 after injection.

#### 2.7.3. Seizure frequency

Seizures were annotated and counted by researchers (J.S., M.D., T.M.) blinded to the treatment. They were defined as a repetitive pattern (> 2 Hz) of high-amplitude EEG spikes (>2x amplitude of background EEG) that lasted a minimum of 7 seconds and were separated by at least 7 seconds from a previous seizure. The seizure frequency was calculated per 30 minutes and normalized to the 2-hour baseline period.

#### 2.7.4. Rotarod

The latency to fall, expressed in seconds, was recorded for each trial on the accelerating rotarod, with a maximum latency of 300 seconds if animals completed the trial without falling. One trial (one animal with vehicle treatment) where the animal turned around on the rotarod and started walking backwards was excluded.

### 2.8. Statistical analysis

Statistical analyses were performed using SPSS Statistics (version 29.0, IBM Corp., USA). For the acute EP recordings, the effects of CPA and cCPA administration were compared between the vehicle, low dose CPA and high dose CPA groups and between the cCPA-light and cCPA-dark groups, respectively. The effects over time on EP parameters and EEG power were evaluated using a restricted maximum likelihood (REML) linear mixed effects model with either fEPSP amplitude, PS amplitude or EEG power as dependent variable, group, time and group by time interaction as fixed factors and animal ID as random factor. The first-order autoregressive model was used as covariance structure to account for repeated measures over time. Fisher’s Least Significant Difference (LSD) test was used for *post-hoc* comparison of timepoints between groups.

For the EEG recordings in the IHKA model, the average seizure frequency and interictal EEG power of the 2-hour period after injection were compared using a REML linear mixed effects model with either seizure frequency or EEG power as dependent variable, treatment condition and treatment period (1^st^, 2^nd^ or 3^rd^ injection) as fixed factors and animal ID as random factor. Compound symmetry was used as covariance structure and Fisher’s LSD for *post-hoc* comparison.

For the data from the rotarod test, the performance on the accelerating rotarod and EEG power were compared between conditions. A REML linear mixed effects model was constructed with either latency to fall or EEG power as dependent variable, treatment condition as fixed factor, animal ID as random factor, using compound symmetry as covariance structure and Fisher’s LSD for *post-hoc* comparison.

For all statistical analyses, a two-tailed *p*-value < 0.05 was set for statistical significance. Group values are reported and plotted as means ± standard error of the mean.

## 3. Results

### 3.1. Effects of A1R activation on hippocampal electrophysiology

We measured hippocampal EPs in healthy mice under anesthesia to confirm that ICV administered cCPA can reach the hippocampus in sufficient levels so that local photo-uncaging of CPA can induce A_1_R signaling and reduce neurotransmission and excitability in hippocampal neurons. First, we validated the expected decrease in EP parameters upon ICV administration of CPA. Then, we examined whether a similar reduction could be achieved using ICV administration of cCPA combined with local illumination.

#### 3.1.1. Dose-dependent suppression of EPs after ICV administration of CPA

Changes in EP parameters and EEG power were studied after ICV injection of two doses of CPA: 0.25 µg (low dose, n = 5) and 1.25 µg (high dose, n = 7). Three out of 7 animals that received the high dose of CPA died within 20 minutes after injection, resulting in 4 complete recordings. Injection of CPA resulted in a dose-dependent decrease in fEPSP and PS amplitudes (**Figure 3 B-C**). The PS was most sensitive to effects of CPA administration, reaching a minimum of 62 ± 13% of baseline levels in the low dose group (n = 5) and a minimum of 22 ± 7% in the high dose group (n = 4) between 30-35 minutes after injection. Respectively, fEPSP amplitudes reached minima of 76 ± 10% and 51 ± 11%. For the PS amplitude, there was a significant effect of the group factor (*F* = 12.085, *p* = 0.001) with both the low and high dose groups significantly differing from the vehicle group starting 10 minutes after injection, and with a significant difference between the two doses starting 20 minutes after injection. For the fEPSP amplitude, the effect of the group factor was near statistically significance (*F* = 3.677, *p* = 0.053) and the high dose group reached a significant difference from vehicle 20 minutes after injection and from the low dose group 30 minutes after injection in *post-hoc* testing. Spectral analysis of hippocampal EEG also displayed a dose-dependent decrease in EEG power (**Fig 3 D-E**). The average total EEG power (1-100 Hz) was reduced with 4.1 ± 1.4 dB from baseline in the low dose group (n = 5) and 6.1 ± 3.2 dB in the high dose group (n = 3) between 10 and 20 minutes post-injection. Mainly the lower frequency bands were affected by CPA administration. LMM analysis revealed a significant effect of group for the delta band (*F* = 9.022, *p* = 0.004) and the theta band (*F* = 5.946, *p* = 0.018), where both doses achieved significant reductions in power compared to vehicle, while the beta band (*F* = 0.858, *p* = 0.452) and gamma band (*F* = 1.144, *p* = 0.358) showed no significant differences.

**Figure 3.**
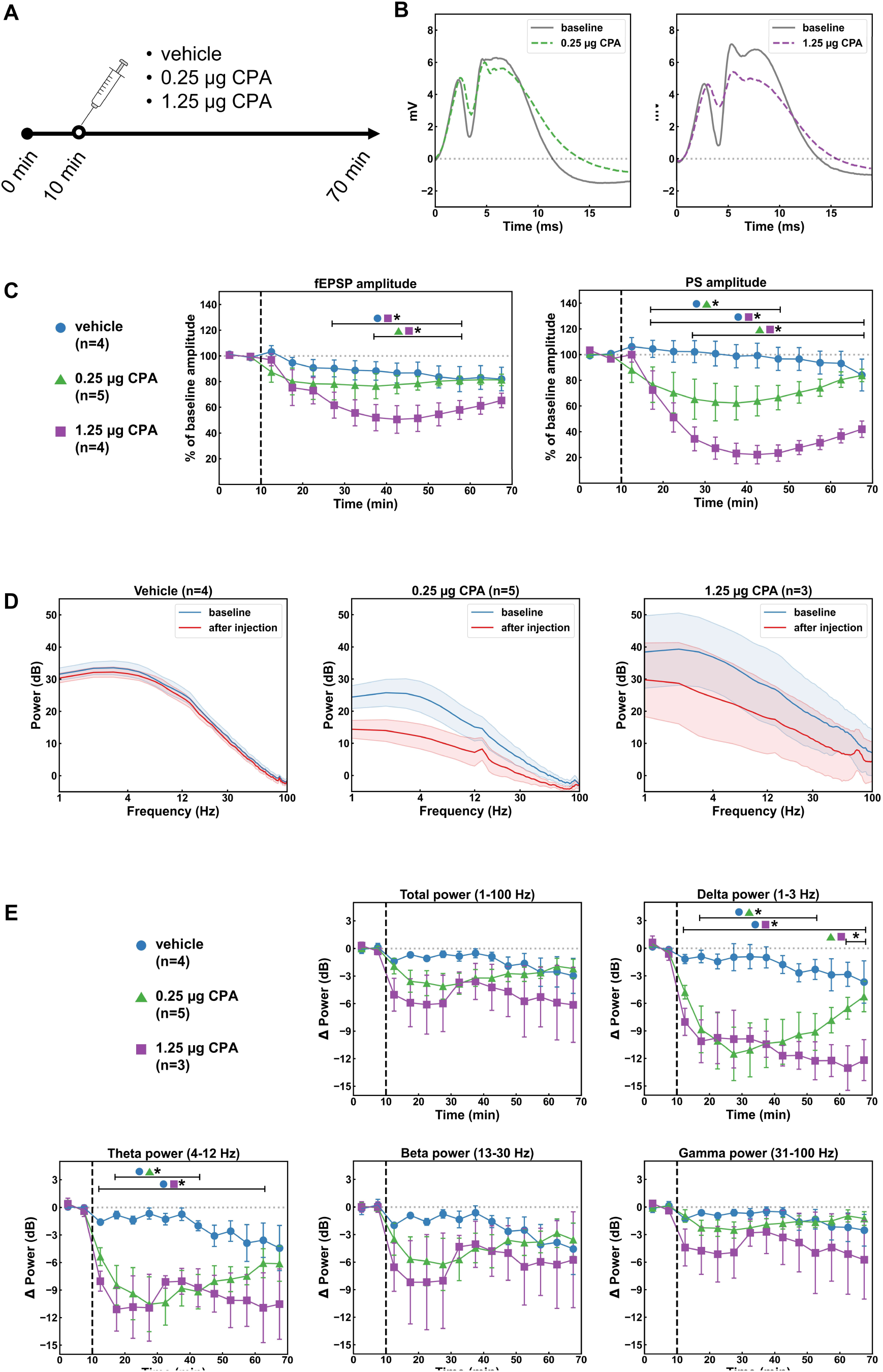
Effects of intracerebroventricular (ICV) injection of CPA on hippocampal evoked potentials (EPs) and electroencephalography (EEG) power. (**A**) Recording protocol: after 10 minutes of baseline EP recording, animals received an ICV injection of 0.25 μg CPA (n = 5), 1.25 μg CPA (n = 4) or vehicle (2.5% DMSO, n = 4) and recordings were continued for 60 minutes. (**B**) Example traces of EPs recorded before (solid line) and 30 minutes after (dashed line) administration of both doses of CPA (left: 0.25 µg CPA; right: 1.25 µg CPA). (**C**) Dose-dependent effects of CPA on field excitatory post-synaptic potential (fEPSP) and population spike (PS) amplitudes over time. Dashed line indicates time of injection. **\*** indicates timepoints where groups (as indicated by the symbols) significantly differ from each other with p < 0.05. (**D**) EEG power spectra before (0-10 min, blue) and after (20-30 min, red) ICV injection of vehicle (n=4, left), 0.25 µg CPA (n=5, middle) and 1.25 µg CPA (n=4, right). (**E**) Change over time in EEG power compared to baseline in the different frequency bands (total power:1-100 Hz, delta power:1-3 Hz, theta power: 4-12 Hz, beta power: 13-30 Hz, gamma power: 31-100 Hz). Dashed line indicates time of injection. **\*** indicates timepoints where groups (as indicated by the symbols) significantly differ from each other with p < 0.05.

#### 3.1.2. Suppression of EPs through photo-uncaging of cCPA

Next, EP recordings were conducted in combination with ICV injection of 25 µg of cCPA. One group of animals received pulsed 405 nm illumination at the DG (cCPA-light, n = 6) and was compared to a control group without illumination (cCPA-dark, n = 7). No animals died after injection of cCPA in either group. In the cCPA-dark group no significant effects of cCPA administration could be detected. However, in the cCPA-light group there was a clear reduction in fEPSP amplitude and PS amplitude (**Figure 4 B-C**). After 20 minutes of pulsed illumination, the PS amplitude and fEPSP amplitude were reduced to minima of 46 ± 15% and 80 ± 9% of baseline, respectively. LMM analysis yielded a significant effect for the group factor for both EP parameters (PS amplitude: *F* = 5.345; *p* = 0.034; fEPSP amplitude: *F* = 7.009, *p* = 0.018). Uncaging of cCPA also caused a small decrease in EEG power (**Fig 4 D-E**). Ten minutes after the start of illumination, average total EEG power (1-100 Hz) was reduced with 3.0 ± 1.1 dB from baseline in the cCPA-light group. The effects were visible across all frequency bands and confirmed by LMM analysis, with significant differences between groups for each band (total power: *F* = 12.432, *p* = 0.002; delta: *F* = 16.177, *p* < 0.001; theta: *F* = 11.559, *p* = 0.002; beta: *F* = 14.693, *p* < 0.001; gamma: *F* = 21.360, *p* < 0.001).

**Figure 4.**
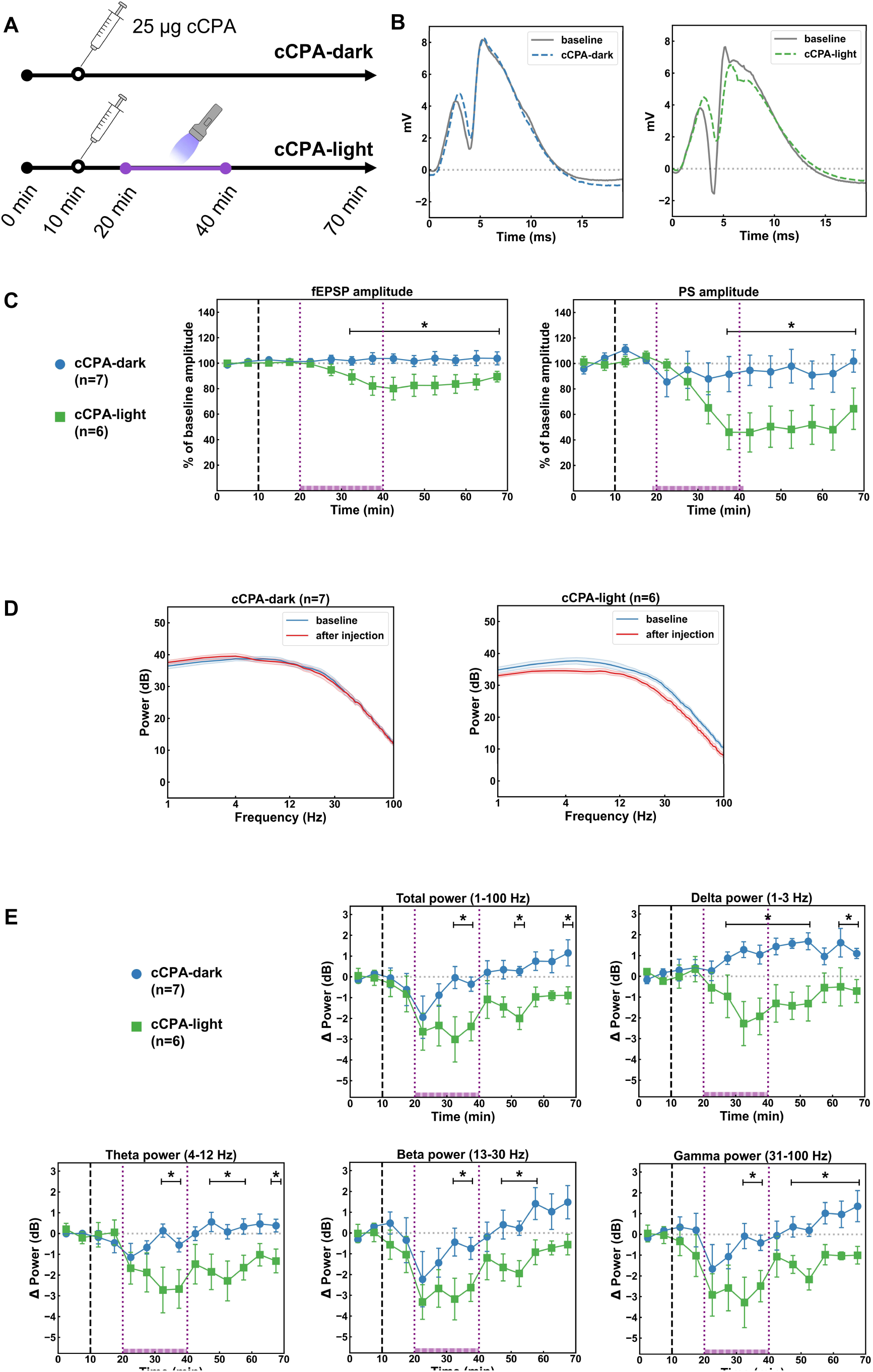
Effects of cCPA photo-uncaging after ICV administration of cCPA on hippocampal evoked potentials (EPs) and electroencephalography (EEG) power. (**A**) Recording protocol: after 10 minutes of baseline EP recording, animals received an ICV injection of 25 µg cCPA and recordings were continued for 60 minutes with (cCPA-light, n=6) and without (cCPA-dark, n=7) 405 nm illumination. Pulsed illumination was started 10 minutes after injection and applied for 20 minutes. (**B**) Example traces of EPs recorded before (solid line) and 30 minutes after (dashed line) administration of cCPA without (left) and with (right) illumination. (**C**) Effects of photo-uncaging of cCPA on field excitatory post-synaptic potential (fEPSP) and population spike (PS) amplitudes over time. Dashed line indicates time of injection, dotted lines indicate start and end of illumination in the cCPA-light group. **\*** indicates timepoints where cCPA-light significantly differs from cCPA-dark with p < 0.05. (**D**) EEG power spectra before (0-10 min, blue) and after (30-40 min, red) ICV injection of cCPA without (left, n=7) and with (right, n=6) illumination. (**E**) Change over time in EEG power compared to baseline in the different frequency bands (total power: 1-100 Hz, delta power: 1-3 Hz, theta power: 4-12 Hz, beta power: 13-30 Hz, gamma power: 31-100 Hz). Dashed line indicates time of injection, dotted lines indicate start and end of illumination in the cCPA-light group. **\*** indicates timepoints where cCPA-light significantly differs from cCPA-dark with p < 0.05.

### 3.2. Suppression of seizures through photopharmacological A1R activation

After confirming a potent reduction in hippocampal neurotransmission induced by local photo-uncaging of cCPA in the acute experiments, we evaluated the seizure-suppressing effects of administering the same ICV dose of cCPA (25 µg) combined with pulsed hippocampal illumination (100 ms pulses of 50 mW 405 nm light at 0.1 Hz) in epileptic IHKA mice.

Based on the presence of clear electrographic seizures during baseline EEG recording, 7 out of 12 animals were selected for inclusion. Animals received 3 treatments in randomized order: vehicle, cCPA-dark and cCPA-light. Administration of cCPA combined with illumination led to a strong decrease in seizure frequency compared to baseline (23± 16% of baseline), where in 3 animals seizures were completely absent during the 2h illumination period (**Figure 5 B-C**). The other conditions had no effect on seizures compared (vehicle: 96 ± 5% of baseline; cCPA-dark: 92 ± 10% of baseline). LMM analysis confirmed that seizure frequency indeed was different between conditions (*F* = 10.133, *p* = 0.005), independent of treatment period (1^st^, 2^nd^ or 3^rd^ injection), with a significantly lower amount of seizures in the cCPA-light condition compared to vehicle (*p* = 0.006) and cCPA-dark (*p* = 0.003) conditions. No statistical differences between vehicle and cCPA-dark were observed. While there was an initial drop in seizure frequency in all 3 conditions immediately after the injections, the course of seizure frequency plotted over time displayed a stable decrease of seizure activity only in the cCPA-light condition during the 2-hour illumination period. In the following 2 hours after illumination, seizure frequencies started increasing back up to around 50% of baseline (**Figure 5 D**). Spectral power analysis of the interictal EEG also showed these differences between conditions, although less pronounced. The average power spectra of the vehicle and cCPA-dark conditions did not differ from baseline, while the average total power (1-100 Hz) in the cCPA-light condition was reduced with 4.1 ± 2.3 dB relative to baseline (**Fig 5 E-F**). The main effects were found in the higher frequency ranges. Power in the theta, beta and gamma bands showed a trend towards reduction in the cCPA-light condition (theta: −5.6 ± 3.7 dB; beta: −5.2 ± 3.3 dB; gamma: −4.1 ± 2.0 dB), but no statistically significant difference was reached (delta: *F* = 0.232, *p* = 0.797; theta: *F* = 1.854, *p* = 0.202; beta: *F* = 1.607, *p* = 0.245; gamma: *F* = 3.302, *p* = 0.076).

**Figure 5.**
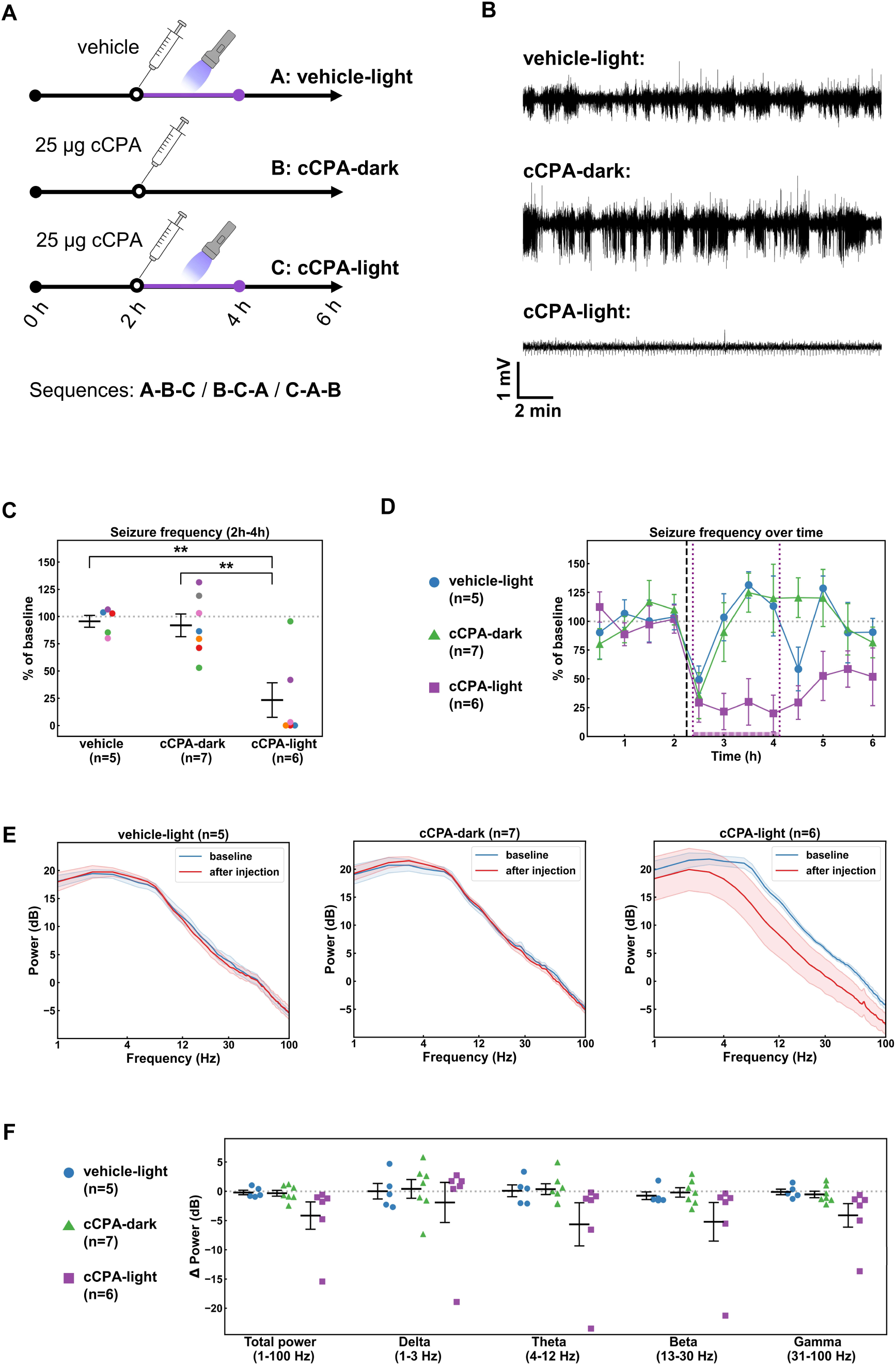
Effects of photo-uncaging of cCPA on epileptic seizures and interictal electroencephalography (EEG). (**A**) Design of the cross-over experiment with cCPA in the intrahippocampal kainic acid (IHKA) model. Animals were randomly assigned to 1 out of 3 sequences to receive 3 treatments: ICV injection of vehicle with 405 nm illumination (vehicle-light), ICV injection of 25 µg cCPA without illumination (cCPA-dark) and 25 µg cCPA with illumination (cCPA-light). Injections were performed after 2 hours of baseline EEG recording, followed by 2 hours of pulsed 405 nm illumination in the vehicle-light and cCPA-light treatments. Two days of wash-out were implemented between treatments. (**B**) Examples of 20-minute EEG segments after administration of the 3 treatments in an animal with complete seizure suppression in the cCPA-light treatment. (**C**) Effects on average seizure frequency during the 2-hour period after injection for the vehicle-light (n=5), cCPA-dark (n=7) and cCPA-light (n=6) treatments. Results of individual animals for each treatment are represented by marks color-coded for each animal. **\*\*** indicates p < 0.01. (**D**) Effects on seizure frequency over time of the 3 treatments, plotted per 30 minutes. Dashed line indicates time of injection, dotted lines indicate start and end of illumination in the vehicle-light and cCPA-light groups. (**E**) Power spectra of the interictal EEG before (blue) and after (red) administration of the 3 treatments. (**F**) Change in interictal EEG power compared to baseline in the different frequency bands (total power:1-100 Hz, delta power:1-3 Hz, theta power: 4-12 Hz, beta power: 13-30 Hz, gamma power: 31-100 Hz). Results of individual animals are represented by marks.

**Figure 6.**
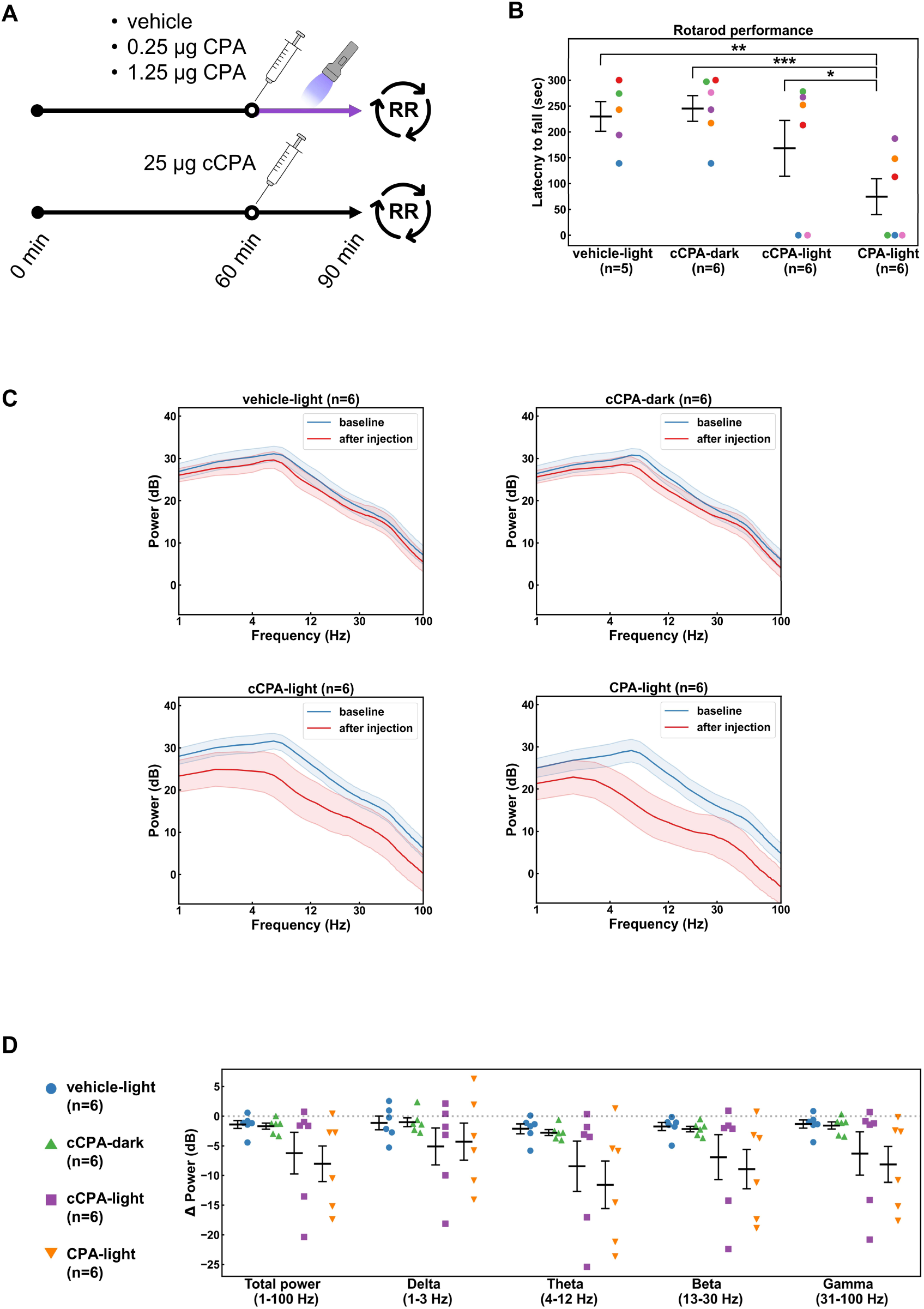
Effects of CPA and cCPA administration on rotarod performance and electroencephalography (EEG) power. (**A**) Protocol of the EEG recordings and rotarod (RR) trials: after 1 hour of baseline EEG recording, animals received an ICV injection of either vehicle with 405 nm illumination (vehicle-light), 25 µg cCPA without illumination (cCPA-dark), 25µg cCPA with illumination (cCPA-light) or 1 µg CPA with illumination (CPA-light). EEG was recorded for 30 minutes after injection, after which animals were placed on the rotarod. (**B**) Effects on rotarod performance 30 minutes after injection for the vehicle-light (n=5), cCPA-dark (n=6), cCPA-light (n=6) and CPA-light (n=6) treatments. Results of individual animals for each treatment are represented by marks color-coded for each animal. **\*** indicates p < 0.05, **\*\*** indicates p < 0.01, **\*\*\*** indicates p < 0.001. (**C**) EEG power spectra before (blue) and after (red) ICV injection for the 4 treatments. (**D**) Change in EEG power compared to baseline in the different frequency bands (total power:1-100 Hz, delta power:1-3 Hz, theta power: 4-12 Hz, beta power: 13-30 Hz, gamma power: 31-100 Hz). Results of individual animals are represented by marks.

Control injections of the photocage DEACM were performed in 3 animals after the cross-over trial. Neither seizure frequency (102 ± 5% of baseline) nor the EEG power spectrum were affected by DEACM compared to baseline (**Supplementary Figure 2**).

### 3.3. Side effects of CPA & cCPA on locomotor activity

To confirm that ICV injection of cCPA was not associated with the typical sedative effects of CPA, we compared the effects of cCPA and CPA administration on locomotor performance in the accelerating rotarod test and on hippocampal EEG. In addition, we evaluated possible sedative effects of photo-uncaging cCPA by local illumination in dentate gyrus.

Intracerebroventricular injection of 1 µg of CPA caused strong sedative effects with a clear reduction in rotarod performance. Half of the animals were too sedated after injection to perform the rotarod trial, while the other half had a notably reduced latency to fall compared to the other conditions, resulting in an overall average latency to fall of 75 ± 35 sec. The latency to fall in cCPA-dark conditions (245 ± 25 sec) was comparable to vehicle-light conditions (230 ± 26 sec). Variable results were seen in cCPA-light conditions: 3 animals performed equally well as in the vehicle-light and cCPA-dark conditions, 1 animal had a decreased latency and 2 animals were too sedated to perform the rotarod trial, resulting in an overall average latency to fall of 168 ± 54 sec. Statistical analysis confirmed a significant effect for the condition factor (*F* = 6.958, *p* = 0.004). Only performance after the CPA treatment was significantly reduced compared to vehicle (*p* = 0.004), cCPA-dark (*p* < 0.001) and cCPA-light (*p* = 0.035) treatments (**Fig 5 B**). Spectral analysis showed that impaired locomotor performances in both the cCPA-light and CPA conditions were associated with reductions in EEG power recorded immediately before the rotarod trials. Similarly to the rotarod results, the average power spectra after vehicle and cCPA-dark treatments did not differ. The cCPA-light conditions were associated with an average reduction of total EEG power (1-100 Hz) of 6.2 ± 3.5 dB relative to baseline, while CPA treatment led to a decrease of 8.0 ± 3.0 dB (**Fig 5 C-D**). The effects on EEG power did not reach statistical significance in the LMM in any of the frequency bands (delta: *F* = 1.093, *p* = 0.383; theta: *F* = 3.142, *p* = 0.57; beta: *F* = 2.596, *p* = 0.091; gamma: *F* = 2.838, *p* = 0.073).

## 4. Discussion

This *in* vivo study is the first to demonstrate that a photocaged A_1_R agonist can be used for light-mediated control of neuronal excitability. Intracerebroventricular administration of cCPA in combination with local 405 nm illumination led to a reduction of both the fEPSP and the PS amplitude, similar to what was observed after direct ICV administration of CPA, indicating successful photo-uncaging of cCPA. Furthermore, administration of cCPA in the absence of illumination did not affect EPs, confirming that the photocage renders the agonist inactive at these effective concentrations. This was further supported by the finding of 43% mortality in mice upon ICV administration of 1.25 µg CPA (3.7 nmol), possibly due to lethal apnea caused by CPA diffusion into the respiratory center in combination with isoflurane anesthesia^45^. Although a more than 10-fold higher molar dose of cCPA (25 µg cCPA = 41.1 nmol) was used, no animals died under anesthesia after injection of cCPA.

These findings are in accordance with our previous study where we demonstrated in an *ex vivo* setting the potential of cCPA to achieve light-controlled activation of adenosine A_1_ receptors, resulting in a reduction of hippocampal excitability due to the well-described pre- and postsynaptic G_i_-mediated effects^36^. Achieving these effects *in vivo* proved more difficult due to the differences in experimental settings between the previously described *ex vivo* MEA recordings and the current *in vivo* EP recordings. In the former, thanks to the 2-dimensional configuration of the brain slices bathed in solution containing cCPA, a single light flash sufficed to elicit changes in EPs. In the current *in vivo* setting, it was more difficult to reach sufficient levels of photo-uncaging in the intact brain. The brain volume that can be illuminated *in vivo* is limited due to high attenuation of light in tissue, which is known as one of the major challenges for the application of photopharmacology *in vivo*^46^. Additionally, only a fraction of the injected cCPA is expected to reach the hippocampus where it is then available for uncaging. Thus, our *in vivo* experiments required relatively high doses of cCPA and repeated light flashes to achieve clear inhibitory effects of A_1_R activation.

Besides validating the photopharmacological modulation of hippocampal excitability with cCPA, the first proof of concept experiments have also been performed to showcase the anticonvulsive potential of caged A_1_R agonists. In recent *ex vivo* experiments with the high-potassium slice model, we were able to suppress epileptiform bursts by controlling hippocampal excitability using cCPA^37^. Now, we also expanded these findings to an *in vivo* epilepsy model, by testing photopharmacological suppression of spontaneous seizures in the IHKA mouse model. Our results with cCPA demonstrate the therapeutic potential of local A_1_R modulation, as uncaging in the sclerotic hippocampus strongly suppresses seizures. In several of the animals, the occurrence of seizures was even completely suppressed during the entire illumination period. This sustained suppression was only achieved when cCPA was combined with local illumination. Control recordings with the photocage DEACM also confirmed that the effects were not due to any potential antiseizure activity of the coumarin derivative^47–49^. Gouder *et al*. were the first to show that administration of an A_1_R agonist can effectively suppress seizures in this mouse model of DRE^18^. In their study, 2-chloro-N^6^-cyclopentyladenosine was administered intraperitoneally together with a non-brain permeable antagonist to demonstrate that the suppression is due to central A_1_R activation. However, this was still affecting receptors throughout the entire CNS. The authors already suggested that local delivery of adenosine into the brain could be effective for the treatment of DRE, as earlier studies had shown that focal injections of adenosine or synthetic analogues had anticonvulsive effects in different epilepsy models^4,6,8,50,51^. Our current study confirms that more locally targeted A_1_R receptor modulation is indeed effective at suppressing seizures in this model for DRE.

As one of the main advantages of this photopharmacological approach would be to avoid side effects by targeting A_1_R activation primarily in the epileptic focus, we investigated whether this would have less sedative effects compared to CPA administration. With the current dose of cCPA and illumination protocol, cCPA treatment led to an overall better performance on the accelerating rotarod compared to CPA administration, which caused a clear impairment in all animals. However, cCPA uncaging was not completely free of side effects as in some animals signs of sedation and decreased locomotor activity did occur. In 2 animals, sedation and suppression of EEG after cCPA-light treatment were on a similar level to that after administration of CPA. The most likely explanation would be that in these mice too much CPA was uncaged and/or leaking into other brain regions. Importantly though, such sedative effects were not witnessed in the prior experiments with the cCPA-light treatment in any of the IHKA mice, indicating that seizure suppression did not rely on such off-target or excessive release of CPA. However, further fine-tuning of our current photopharmacological protocol will be required in future work to minimize the occurrence of such side effects.

Throughout the various experiments, we also examined effects on hippocampal EEG. Overall, A_1_R activation either with CPA or photo-uncaging of cCPA caused a decrease in EEG power. Due to the difference in experimental settings, the effects did vary between experiments. In healthy anesthetized mice, ICV injection of CPA caused a larger decrease in EEG power compared to cCPA uncaging and the effects of CPA were most pronounced in the delta and theta band frequency ranges. This could point towards more widespread effects elicited by CPA administration compared to cCPA. In epileptic animals, photo-uncaging of cCPA also caused a small decrease in interictal EEG power. In these EEG recordings the higher frequencies were affected most significantly, again suggesting a mostly local effect. In the healthy animals that were tested on the rotarod, there was a large overall decrease in EEG power upon CPA administration, but so also in the two mice that were sedated in the cCPA-light conditions.

This study provides a proof of concept for *in vivo* light-controlled activation of A_1_R signaling with our novel caged A_1_R agonist. There are several limitations to this study and multiple important questions remain that will require further study to develop this photopharmacological approach beyond a proof of concept. One of the most important limitations is the lack of knowledge about the pharmacokinetics of cCPA in the current experiments, both before and after uncaging. Compounds were injected into the lateral ventricle contralateral to the site of EP/EEG recording and illumination. This route of administration is a relatively effective way to ensure delivery of compounds to the brain, especially in periventricular regions such as the hippocampus^52^. However, the actual concentration of cCPA that reaches the hippocampus and that would be available for uncaging is unknown. This also makes it difficult to estimate the exact amount of cCPA that was released in our experiments. Besides the question of how much cCPA is released, further detailed investigations will also need to address the temporal and spatial resolution of uncaging, investigating the extent of CPA diffusion after uncaging and the duration of activity of the uncaged compound. Adenosine analogues are more resistant to metabolization and half-life values of CPA determined in the blood of rats have been reported to be 24 minutes *in vitro* and 7 minutes *in vivo*^53,54^. It could thus be possible that after local photo-uncaging, there is enough time for the released CPA to diffuse outside of the targeted region.

Here, our photopharmacological approach was executed in acute experiments. Since epilepsy is a chronic disorder, the aim is to further develop this approach into a chronic treatment option. Others have investigated different strategies for chronic local delivery of adenosine, but this involved a continuous slow infusion or release of adenosine^55^. Apart from the possibility to localize the activity of a drug, photopharmacology offers the advantage of temporal control. This would allow to modulate A_1_Rs only when required, avoiding sustained receptor activation which could induce receptor desensitization and reduce efficacy. Photopharmacological modulation could be well-suited for integration with a closed-loop system, where interventions are put under automatic control of a detection system monitoring physiological parameters of interest^56–58^. This concept was investigated in our lab in an *ex vivo* setting^36,37^. Caged CPA was used together with feedback-controlled light delivery to keep neurotransmission, monitored via fEPSP amplitude, to a predefined level. Similarly, cCPA could be used to stabilize the level of excitability and prevent seizures *in vivo*, either through monitoring of EP amplitudes or through EEG registration combined with seizure prediction algorithms. This would allow for a more titrated release of CPA, possibly further decreasing the chance of sedative effects.

In conclusion, with this work we presented the first application of a photocaged A_1_R agonist *in vivo*. We demonstrated that this caged agonist is inactive upon ICV administration and that it is possible to photo-uncage the agonist with local illumination in the hippocampus of mice, achieving inhibitory effects on hippocampal neurotransmission and significant suppression of spontaneous seizures in the IHKA mouse model for DRE. This proof of concept study showcases the therapeutic potential of photopharmacology, as it allows us to take advantage of the potent anticonvulsive effects of the adenosinergic system in a way that can be further developed to achieve minimal side effects through the high spatiotemporal control afforded by light.

## CRediT authorship contribution statement

**Jeroen Spanoghe:** Conceptualization, Methodology, Formal analysis, Investigation, Writing – Original Draft, Visualization. **Paul Boon:** Conceptualization, Supervision, Project administration, Funding acquisition. **Marijke Vergaelen:** Conceptualization, Methodology, Writing – Review & Editing. **Maren De Colvenaer:** Investigation, Formal analysis. **Tina Mariman:** Investigation, Formal analysis. **Kristl Vonck:** Conceptualization, Supervision, Funding acquisition, Writing – Review & Editing. **Evelien Carrette:** Conceptualization, Project administration, Writing – Review & Editing. **Wytse Wadman:** Conceptualization, Software, Writing – Review & Editing. **Erine Craey:** Conceptualization, Methodology, Writing – Review & Editing. **Lars E. Larsen:** Conceptualization, Software, Formal analysis. **Mathieu Sprengers:** Conceptualization, Writing – Review & Editing. **Jeroen Missinne:** Resources, Writing – Review & Editing. **Serge Van Calenbergh:** Resources, Methodology, Writing – Review & Editing. **Dimitri De Bundel:** Conceptualization, Funding acquisition, Writing – Review & Editing. **Ilse Smolders:** Conceptualization, Funding Acquisition, Writing – Review & Editing. **Robrecht Raedt:** Conceptualization, Writing – Review & Editing, Supervision, Project administration, Funding Acquisition.

## Acknowledgements

The authors thank Robin Lefevere for assisting with part of the experiments. Figures were partially created with Biorender.com.

## Supplementary figures

**Supplementary Figure 1.**
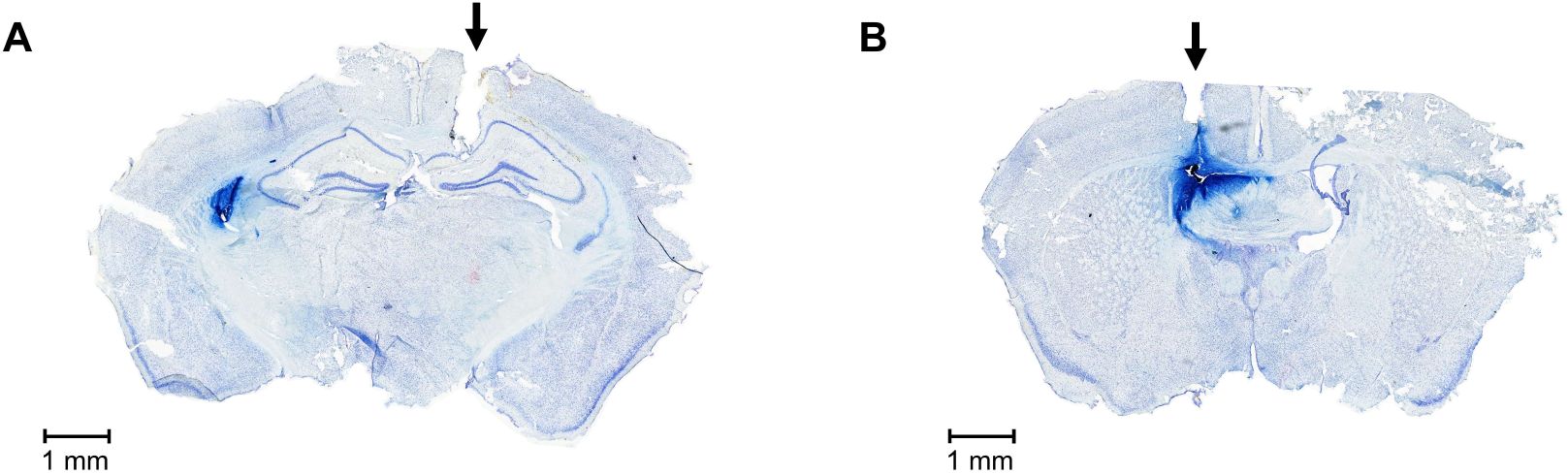
Representative histological images of coronal sections of the mouse brain after intracerebroventricular injection of Evan’s Blue dye and cresyl-violet staining. Arrows indicate the tracts verifying the correct placement of (**A**) the optrode just above the dentate gyrus of the right hippocampus and (**B**) the guide cannula above the left lateral ventricle. Dark blue staining by the dye of the walls of the lateral ventricle shows that injections through the guide cannula successfully targeted the ventricle.

**Supplementary Figure 2.**
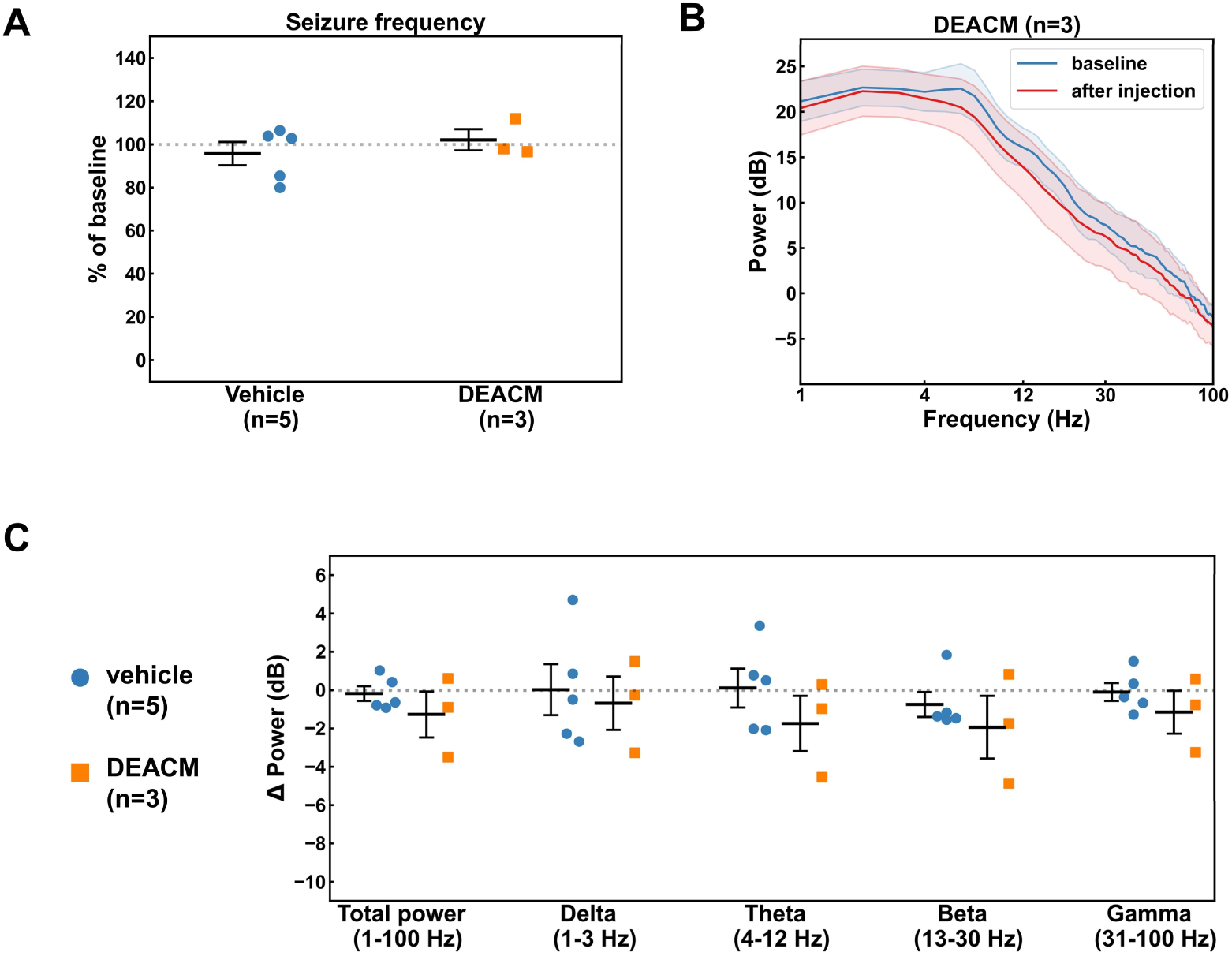
Effects of the DEACM photocage on epileptic seizures and interictal electroencephalography (EEG). (**A**) Effects on average seizure frequency during the 2-hour period after injection of the DEACM solution (n=3) compared to vehicle-light treatment (n=5). (**B**) Power spectra of the interictal EEG before (blue) and after (red) DEACM administration. (**C**) Change in interictal EEG power compared to baseline after DEACM administration compared to vehicle-light treatment in the different frequency bands (total power:1-100 Hz, delta power:1-3 Hz, theta power: 4-12 Hz, beta power: 13-30 Hz, gamma power: 31-100 Hz).

